# Uni-Fold Symmetry: Harnessing Symmetry in Folding Large Protein Complexes

**DOI:** 10.1101/2022.08.30.505833

**Authors:** Ziyao Li, Shuwen Yang, Xuyang Liu, Weijie Chen, Han Wen, Fan Shen, Guolin Ke, Linfeng Zhang

## Abstract

Deep folding models have revolutionized the conventional methods of protein complex prediction. However, applying them to large protein oligomers is not easy. These models generally require copying the sequences of identical subunits to capture the in-between relationships. Accordingly, the scales of target protein complexes are strictly limited due to the cubic complexity of these models. To address this issue, we propose UF-Symmetry (Uni-Fold Symmetry), which is extricated from the need of sequence copying via harnessing the intrinsic symmetry of large protein oligomers. Taking the sequences of the asymmetric unit (AU) and a pre-specified symmetry group, UF-Symmetry learns to fold the AU and to assemble the complex structure in an end-to-end manner. By reducing the input scales from entire assemblies to AUs, UF-Symmetry allows to predict much larger assemblies with significant acceleration: for a complex of 4-fold cyclic symmetry (C4) and AU size of 512, UF-Symmetry achieves approximately 20 times acceleration to current methods. On a benchmark of recently released PDB multimers, UF-Symmetry approximately halves the failure rate of current methods and achieves approaching accuracy on commonly successful cases.

## 1 Introduction

Understanding the structures of proteins and their complexes is a key preliminary to life science research and structure-based drug discovery. *Deep folding models*, which predict unknown protein structures via learning from solved ones with neural networks, are now revolutionizing the domain. Extending the success of AlphaFold [1], folding models for protein multimers [2, 3] have now significantly improved the accuracy of conventional docking methods [4, 5].

A fascinating phenomenon of protein complexes is that a majority of them function as large and symmetric oligomers^2^. As is shown in Figure 1, about 72.1% multimeric structures in the Protein Data Bank (PDB) [6] are globally symmetric. Symmetric proteins are evolutionarily preferred due to functional, genetic, and physicochemical needs [7]. For instance, *ion channels* are typically formed as assemblies with a circular arrangement of identical (or homologous) proteins closely packed around a water-filled pore through the plane of the lipid bilayer. Solving the structures of these symmetric assemblies yields great importance to understanding the mechanisms of relevant biological activities.

**Figure 1:**
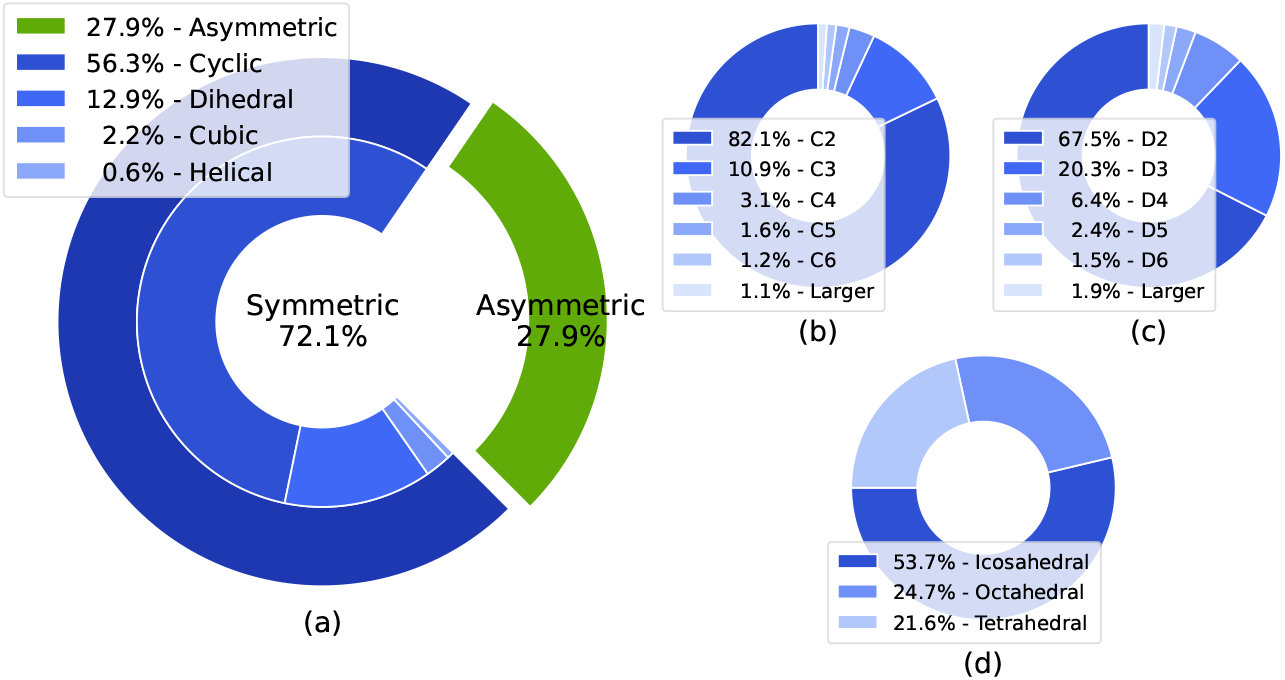
Statistics of symmetric structures in PDB multimers. **(a)** Proportions of different symmetry types in all PDB multimers. **(b) (c) (d)** Proportions of different cyclic, dihedral and cubic symmetries, respectively. Detailed explanations of this figure are in Section A.

Despite the remarkable accuracy of current deep folding models on small complexes, applying them to large oligomers is not easy, specifically because of the combination of two facts: i) these models generally require copying the sequences of the repeating subunits to capture the in-between connections, typically with the self-attention mechanism [8]; ii) the complexities of these models are generally cubic to the total length of the input sequences. Taking Uni-Fold Multimer [3], one of the fastest and most memory-efficient deep folding models as an example, an NVIDIA A100 GPU with 40GB memory allows the inference of up to ∼ 2,100 residues^3^ due to memory limitation. For large oligomers, the input scales can easily go beyond this limit.

Fortunately, harnessing the intrinsic symmetry of large oligomers may help to solve the problem: as the *asymmetric units* (AUs)^4^ are structurally identical in an oligomer, one can predict the AU structure and then assemble the entire complex. This saves the effort of repeatedly predicting the AUs. Moreover, one can generate oligomer structures with different symmetries (or stoichiometries) by querying the model with different symmetry groups.

Following this motivation, we propose **UF-Symmetry (Uni-Fold Symmetry)**, which predicts protein complex structures based on the AU sequences and a pre-specified symmetry group. UF-Symmetry learns to *fold* the AU and to *assemble* the oligomer in an end-to-end manner. To briefly summarize, the model modifies the pipeline of Uni-Fold Multimer to fold the AUs with a newly introduced *pseudo residue*, which encodes the given symmetry type and learns to correctly pose the AU in a standard frame. Defined by the input symmetry group, a group of standard symmetry operations is then applied to the AU to compose the final assembly. To efficiently supervise the process, we further reduce the complexity of the structure losses by merging the identical intra-AU terms. This also allows the model *not* to explicitly generate the assembly structures during training.

By harnessing symmetry, UF-Symmetry reduces the model complexity from *O*(*K*^3^*N* ^3^) to *O*(*N* ^3^), where *K* is the number of AUs in the assembly and *N* is the AU size. Notably, this new complexity is invariant to the number of AUs in the assembly, which not only allows the model to be applied on larger oligomers, but also significantly accelerates the inference process. We also evaluated the accuracies of UF-Symmetry and baselines on 416 recently released protein multimers in PDB. AlphaFold-Multimer and Uni-Fold Multimer failed on 11.1% cases due to out-of-memory (OOM) errors, while UF-Symmetry reduces this rate to 5.8%. In commonly successful cases, UF-Symmetry displayed approaching accuracy, falling behind AlphaFold-Multimer of ∼ 2% TM-Score. We reckon the performance drop as a necessary trade-off for larger model capacity. Further analysis shows that despite the drop of average performance, UF-Symmetry is yet more competent on a considerable proportion of specific cases. Therefore, using UF-Symmetry in complement to Uni-Fold Multimer well balances the model accuracy and flexibility.

## 2 Preliminaries

### Protein Symmetry

All amino acids but glycine have chirality over the *α* carbon, and nature dominantly prefers L-amino acids to D-ones. Consequently, natural proteins adopt only enantiomorphic symmetries and disallow mirror and inversion ones. In this paper, the Schönflies symbols are used to denote symmetry groups, where C*x* is for *x*-fold cyclic groups, D*x* for *x*-fold dihedral groups, T, O, I for tetrahedral, octahedral and icosahedral groups respectively, and H for helical structures. In UF-Symmetry, we focus on proteins with finite symmetry groups (C, D, I, O, T), and leave helical structures as future work.

### Notations

We use ***X*** = (***x***_1_,, ***x***_*n*_)^*T*^ ∈ ℝ^*n*×3^ to denote the structure of an AU, where ***x***s are the 3D coordinates of all heavy atoms (or C_*α*_ atoms, defined by the context). 𝒮 = ***X***_*i*_ denotes an assembly as a set of AU structures. A symmetry operation *O* = (***R, t***) is a transformation on the 3D euclidean space, consisting of a rotation matrix ***R*** ∈ SO(3) and a translation vector ***t*** ∈ ℝ^3^. The operation can be applied to a point ***x***, an AU structure ***X***, or composed with another operation *O*^′^ = (***R***^′^, ***t***^′^) as

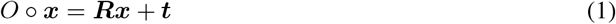

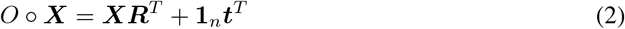

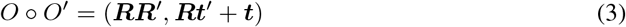

A symmetry group is a set of symmetry operations 𝒢 = {*O*_*i*_} which is closed under composition. An assembly *conforms to* a symmetry group 𝒢 if for any ***X*** ∈ 𝒮 and *O* ∈ 𝒢, one has *O* ○ ***X*** ∈ 𝒮.

### Deep Folding Models

The backbone model of UF-Symmetry is originally inspired by AlphaFold [1]. Taking the amino acid sequence, homologous sequences and solved homologous structures as inputs, AlphaFold directly predicts the three-dimensional coordinates of all atoms in a protein. In AlphaFold, an *Evoformer* encodes the inputs with the attention mechanism, and a *structure module* decodes the local frames and torsion angles of the protein residues. The *Frame Aligned Point Error* (FAPE) loss is used to supervise the output structure by averaging the structure errors superposed by each residue. The *violation loss* is used to encourage the correct peptide bond geometry and to punish steric clashes. AlphaFold-Multimer [2] and Uni-Fold Multimer [3] further extended AlphaFold to protein complexes by slightly modifying the system and retraining it on multimeric protein structures in PDB. The backbones of these models are hardly modified, whereas an MSA pairing strategy is introduced to build cross-chain genetics. An additional *chain center-of-mass* loss is also introduced to teach the model the global placement of the chains.

## 3 Method

Figure 2 shows the pipeline of UF-Symmetry. Given the input AU sequences and a symmetry type, the model first conduct the homology search on sequence and structure databases. The obtained MSAs and templates are then padded with the pseudo residue and encoded by the Evoformer. The structure module then decodes the local frames of the AU residues and transforms them into the global frame learned by the pseudo residue. In inference, a group of standard symmetry operations are applied to the AU structure to form the final assembly. In training, reduced structure-related losses (symmetry losses) are directly calculated with the AU structure and the standard symmetry group.

**Figure 2:**
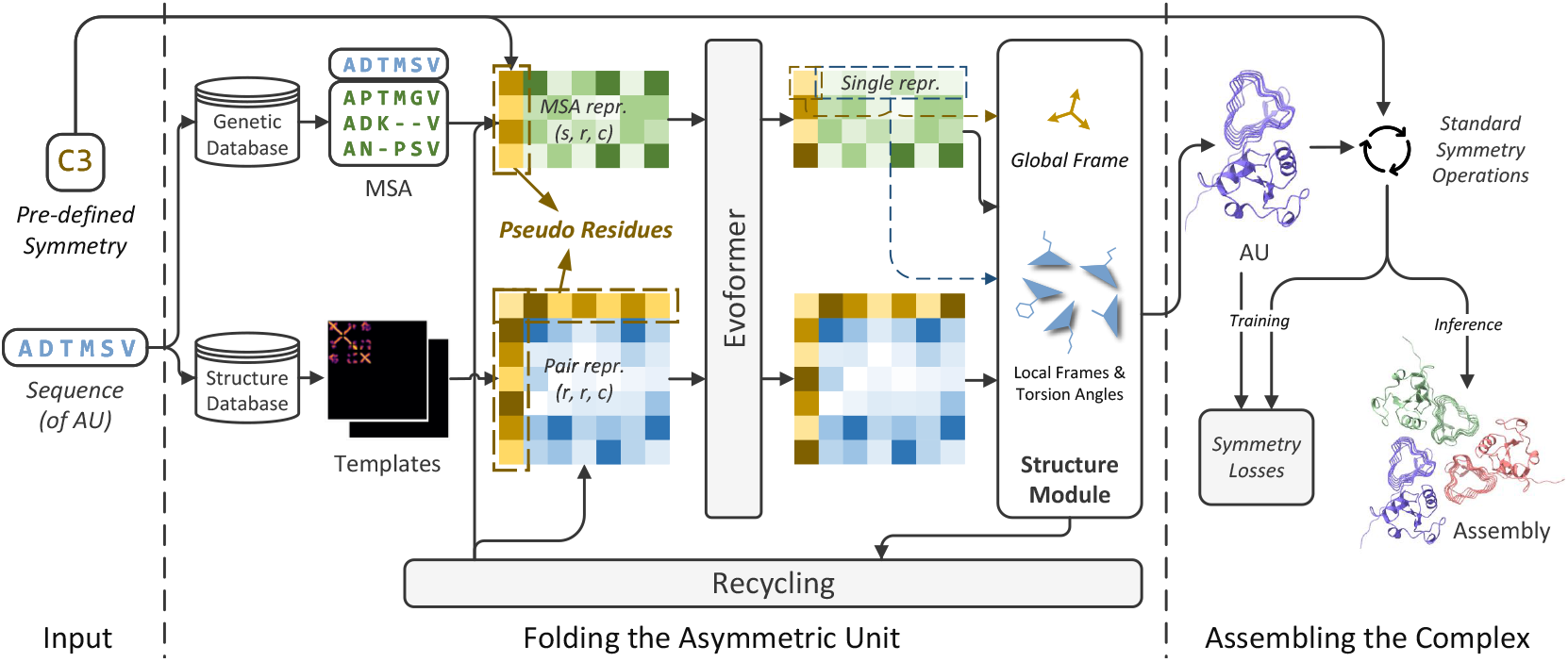
The pipeline of UF-Symmetry. dock

### 3.1 Folding the Asymmetric Unit

The AU prediction process in UF-Symmetry is modified from the structure prediction process in Uni-Fold Multimer. Unless otherwise specified, the implementation details are the same.

UF-Symmetry introduces a pseudo residue to encode the query symmetry type and to learn a global frame for the AU. The pseudo residue is a placeholder symbol that joins the calculations of the Evoformer and the structure module equally as a real residue. Intuitively, the pseudo residue plays a similar role to the *classification token* ([CLS]) in pre-trained language models such as BERT [9], which collects the global information of the sequences via the self-attention mechanism [8]. This is particularly important in UF-Symmetry, because we are not only interested in how the AU is folded, but also how it is globally posed and repeated to form the oligomer.

The initial features of the pseudo residue consists of the one-hot encoding of the query symmetry type (among [C1, C, D, I, O, T]), and the cosine and sine values of the rotation angles of cyclic and dihedral symmetries (e.g. 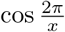 and 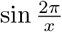 for C*x*). For C1, I, O, and T, the rotation angles are defined as 0. A 4-block residual network transforms these features into the same dimension as the representations of real residues, which is then padded to the front of both the input sequences and each MSA row. For pair representations, the pseudo residue is processed as a real residue, except that we do not use relative position embedding between the pseudo residue and the real residues.

As the pseudo residue joins the calculation of the structure module, a frame, which we name as the *global frame*, is generated for the pseudo residue. When decoding the atom coordinates of the AU, UF-Symmetry transforms the local frames of residues into this global frame. Denoted in formula,

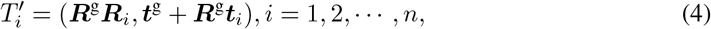

where *T*_*i*_ = (***R***_*i*_, ***t***_*i*_) and *T* ^g^ = (***R***^g^, ***t***^g^) are the real residue frames and the global frame, respectively. 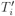 are the final local frames of residues. No torsion angles and side-chain frames are generated for the pseudo residue. In the recycling process, the translation of the global frame is recycled as the C_*β*_ coordinates of the pseudo residue.

### 3.2 Assembling the Complex

In order to generate the entire complex, we first define the standard symmetry groups for different symmetry types. All operations in the standard symmetry groups have zero translations (***t*** = **0**), and the identical operation (*O*_*I*_) is always included in the groups. For cyclic symmetries (C*x*), we use the *z* axis (with unit vector (0, 0, 1)) as the rotation axis, and include the corresponding *x*-fold rotation group, i.e. operators that rotates the AU for 2*kπ/x, k* = 0, 1, …, *x* − 1. For dihedral symmetries (D*x*), the group is the multiplication of the standard C*x* group and a 2-fold rotation group over the *y* axis ((0, 1, 0)). For the cubic symmetries, the standard operations are of more complicated forms, and are described in Section B. For asymmetric structures (C1), the group is {*O*_*I*_ }.

The assembling process does not introduce additional learnable parameters. By presuming that the AU (with structure ***X***_1_) is properly posed in the standard frame associated with the standard symmetry groups, it simply applies all operations in the standard group of the given symmetry type to the AU structure, i.e. 𝒢 = {*O*_*i*_ ○ ***X***_1_ | *O*_*i*_ ∈ 𝒢}. Accordingly, the generated structure necessarily conforms to the given symmetry: for all *O*_*i*_ ∈ 𝒢 and ***X***_*j*_ ∈ 𝒮, one has

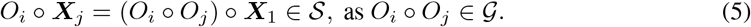

In inference, the assembling process generally takes milliseconds, negligible to the total time. Accordingly, the inference time of UF-Symmetry stays invariant to the size of the symmetry group.

### 3.3 Symmetry Losses

As the assembling process is differentiable to the AU structure, directly applying the losses of Uni-Fold Multimer on UF-Symmetry is feasible. However, the complexity of the Frame Aligned Point Error (FAPE) loss is squared to the total number of residues, and thus the full calculation of the losses also leads to insufficient memory for large-scale complexes. To address this issue, we reduce the structure-based losses to lower computational complexity without explicitly using the assembly structure. Intuitively, the reduction is achieved by merging the identical loss terms of AU pairs that enjoys the same symmetry operation, as is introduced by Theorem 1:

#### Theorem 1

*For any (finite) symmetric assembly* 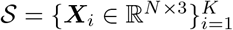 *and its symmetry group* 𝒢, *assuming* ℱ : 𝒮 ×𝒮 → ℝ *is a real-valued function of a pair of AUs and is invariant to the symmetry operations, i*.*e*. ℱ (*O* ○ ***X***_*i*_, *O* ○ ***X***_*j*_) = ℱ (***X***_*i*_, ***X***_*j*_) *for all O* ∈ 𝒢, *one has*

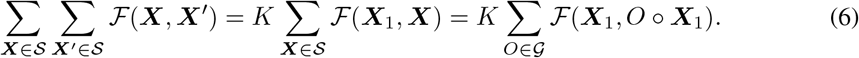

Accordingly, the complexity of loss calculation is reduced from *O*(*K*^2^*N* ^2^) to *O*(*KN* ^2^).

Losses that require the assembly structure include the FAPE loss, the violation loss, and the chain centre-of-mass loss. For FAPE loss, given the symmetry group 𝒮, we calculate the whole loss as

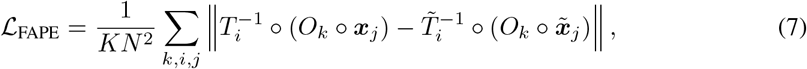

where *T*, ***x*** are the local frames and coordinates of predicted residues, and 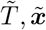 those of ground-truth. The index *k* is iterated over 𝒢, and *i, j* over the residues inside the AU. For the violation loss, we reduce the clash terms of atom pairs between different AUs following the same idea:

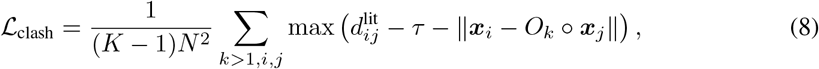

where 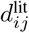 is the clashing distance between the atom types of *i* and *j*, and *τ* is a tolerance factor. We do not alter other terms of the loss calculated inside the AU. The between-AU term of the chain centre-of-mass loss is modified similarly as

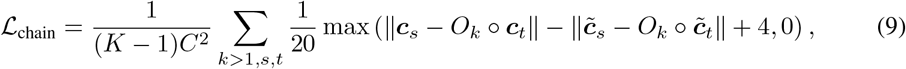

where indices *s, t* are iterated over all chains (totally *C* chains) in the AU, and ***c***s are the coordinates of the chain centers. Notably, applying these losses requires first transforming the ground-truth (*T* and 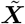) into the standard frame. Choosing any one of the AUs among the ground-truth is equivalent.

The proof of Theorem 1 and the equivalency of Equations (7), (8) and (9) are in Section C.

### 3.4 Symmetry Augmentation

In order to encourage robustness of the model to any given symmetry group, we propose *symmetry augmentation* as a training strategy of UF-Symmetry. Specifically, with a certain probability, we replace the annotated symmetry group of the target structures with one of its subgroups. Taking symmetry group C6 as an example, we may use the subgroup C3 (chosen from C1, C2, and C3) and merge two adjacent AUs into a new one. The probability of using symmetry augmentation is 0.382, and the subgroups are uniformly sampled from all possible ones.

## 4 Results

### 4.1 Training UF-Symmetry

#### Data

We trained UF-Symmetry basically following Uni-Fold Multimer. The training set of UF-Symmetry is the same as Uni-Fold Multimer. Specifically, we collected a total of 181,727 PDB structures released before January 16th, 2022, including both symmetric and asymmetric ones. We used the annotated symmetry groups of the RCSB website^5^, extracted the AUs, transformed them into the standard frame, and used them as labels. Notably, we only extracted the AUs of cyclic structures, while the extension to dihedral and cubic cases is natural. We used a rule-based algorithm in combination with libmsym [10] to extract and transform the AUs. For dihedral and cubic structures, as well as approximately 3% cyclic structures that we failed to extract or transform the AUs, we regarded them as asymmetric ones. We used the identical pipeline to search MSAs and templates to Uni-Fold Multimer, including the database versions and hyper-parameters. We also used the same self-distillation datasets, which contained approximately 360,000 monomeric structures.

#### Recipe

We trained UF-Symmetry on 128 NVIDIA A100 GPUs with 40GB memory, each containing one training sample. The sequence(s) of AUs were cropped to 320 residues using the *contiguous cropping* strategy of [2]. We trained the model for 80,000 steps, which took us approximately one week. We tried to finetune the model under several configurations, while the improvements were minor or negative. To feed the model with more symmetric samples, we decrease the proportion of self-distillation samples from 0.5 to 0.382. Other settings were the same as Uni-Fold Multimer.

### 4.2 Capacity and Efficiency Benchmark

We compared the capacity to input sizes and the inference efficiency of UF-Symmetry and AlphaFold-Multimer. We used the implementation of AlphaFold in the Uni-Fold repository^6^, which was proved as faster and more memory-efficient. Note that as Uni-Fold Multimer and AlphaFold-Multimer share almost identical architecture, these results also apply to Uni-Fold Multimer.

Figure 3 shows the results. We report the performances under different AU and symmetry group sizes. As AlphaFold-Multimer requires copying the sequences of AUs, the maximum AU size the model can handle is in inverse proportion to the size of the symmetry group. In addition, the inference time of AlphaFold-Multimer is also cubic to the size of the symmetry group. By contrast, UF-Symmetry has invariant capacity and inference time to different symmetry group sizes, yielding significant improvements in larger oligomers. For example, given the AU size of 512 and the symmetry group C4, the inference speed of UF-Symmetry is approximately 19.4 times to AlphaFold-Multimer.

**Figure 3:**
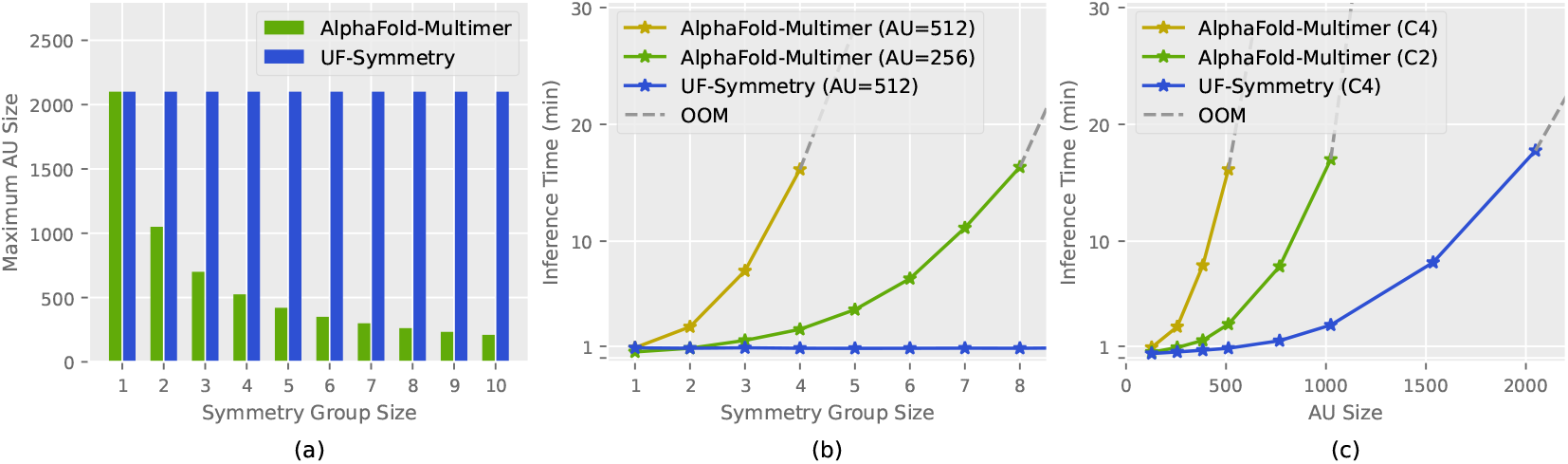
Capacity and inference efficiency of UF-Symmetry and AlphaFold-Multimer on an NVIDIA A100 GPU with 40GB memory. **(a)** The maximum AU sizes the models support given different symmetry group sizes. **(b) (c)** The inference time of models given different AU and symmetry group sizes. Bracketed notes beside the model names describe the fixed AU sizes or symmetry groups. Gray dashes indicate that the next data point is infeasible due to memory limitation.

### 4.3 Accuracy Benchmark

#### Data, Metrics, and Baselines

We evaluated UF-Symmetry and baseline models on recently released multimeric structures in PDB, focusing on the performances of asymmetric and cyclic structures. We collected a total of 1,181 PDB structures released between January 17th and July 14th, 2022, and kept the asymmetric and cyclic multimeric structures, i.e. the structures in C*x* symmetry groups with at least 2 chains. We further filtered the structures such that all structures had resolutions below 3.5 Å and had at least one chain with less than 40% template identity, yielding a total of 416 samples. Statistics of the test samples are in Figure 4. Following Uni-Fold Multimer, we report the C_*α*_-RMSD and TM-Scores of protein complexes. As the C_*α*_-RMSD of entire complexes suffered from large outliers, we truncated the larger values to 30 Å, which we named as C_*α*_-RMSD_30_. These scores were calculated by aligning the entire complexes and averaging the metrics among all residues. We fed UF-Symmetry with the annotated symmetry groups from RCSB. We tested the performances of Uni-Fold Multimer, and all 5 models of AlphaFold-Multimer. We also tested UF-Symmetry (C1) by feeding UF-Symmetry with C1 symmetry and merging all AUs into one.

**Figure 4:**
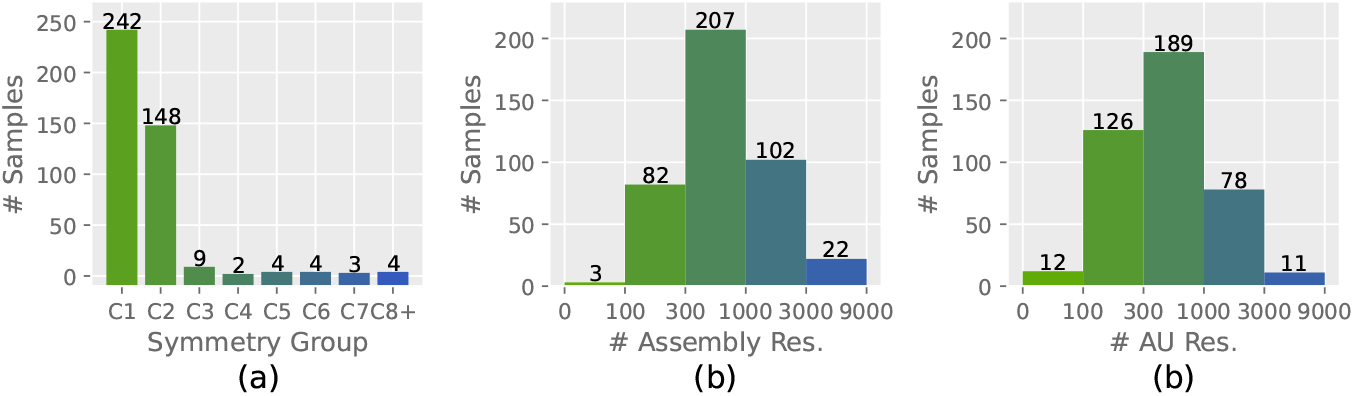
Statistics of the test dataset. **(a)** Number of samples of different symmetry groups. **(b) (c)** Histogram of samples with different assembly / AU sizes.

#### Results

Table 1 shows the evaluation results. We report the number of failed cases due to memory limitations, and the averaged accuracy metrics on commonly successful cases (370 samples). By harnessing symmetry, UF-Symmetry succeeded on 22 more samples compared with AlphaFold-Multimer and Uni-Fold Multimer, which approximately halved the failure rate. On commonly successful samples, UF-Symmetry displayed approaching accuracy, with an approximately 2% drop on the TM-Score compared with AlphaFold-Multimer, and 6% that with Uni-Fold Multimer. Nevertheless, the performances of UF-Symmetry (C1) are better than UF-Symmetry and equivalent to AlphaFold-Multimer, slightly below Uni-Fold Multimer.

**Table 1:** Number of failed cases and prediction accuracies on commonly successful cases of UF-Symmetry and other baselines.

| Model | Failed Cases | $C_{\alpha}$-RMSD<sub>30</sub> (Å) | TM-Score |
| --- | --- | --- | --- |
| AlphaFold-Multimer | 11.1% (46) | 8.40 | 0.757 |
|  |  | 8.26 | 0.755 |
|  |  | 8.62 | 0.744 |
|  |  | 8.19 | 0.758 |
|  |  | 8.30 | 0.756 |
| Uni-Fold Multimer | 11.1% (46) | **6.98** | **0.791** |
| UF-Symmetry (C1) | 9.4% (39) | 8.77 | 0.760 |
| UF-Symmetry | **5.8% (24)** | 9.74 | 0.733 |

The comparison between UF-Symmetry and UF-Symmetry (C1) may help to reveal the reason of the performance drop: as we do not copy the AU sequences, the intra-AU connections cannot be captured as well as those in UF-Symmetry (C1) as well as other multimer models. This could be considered as a trade-off between complexity and accuracy. Meanwhile, the smaller cropping sizes and the lack of finetuning may account for the accuracy drop from Uni-Fold Multimer to UF-Symmetry (C1).

More detailed accuracy results are shown in Figure 5, in which we draw both TM-Score and C_*α*_-RMSD_30_ as scatter plots. Although the averaged accuracies of UF-Symmetry drops, it still attains competitive performances on specific cases (indicated with the blue dots). Therefore, in real scenarios, one may use UF-Symmetry in complement to Uni-Fold Multimer or UF-Symmetry (C1) to achieve the best balance between flexibility and accuracy.

**Figure 5:**
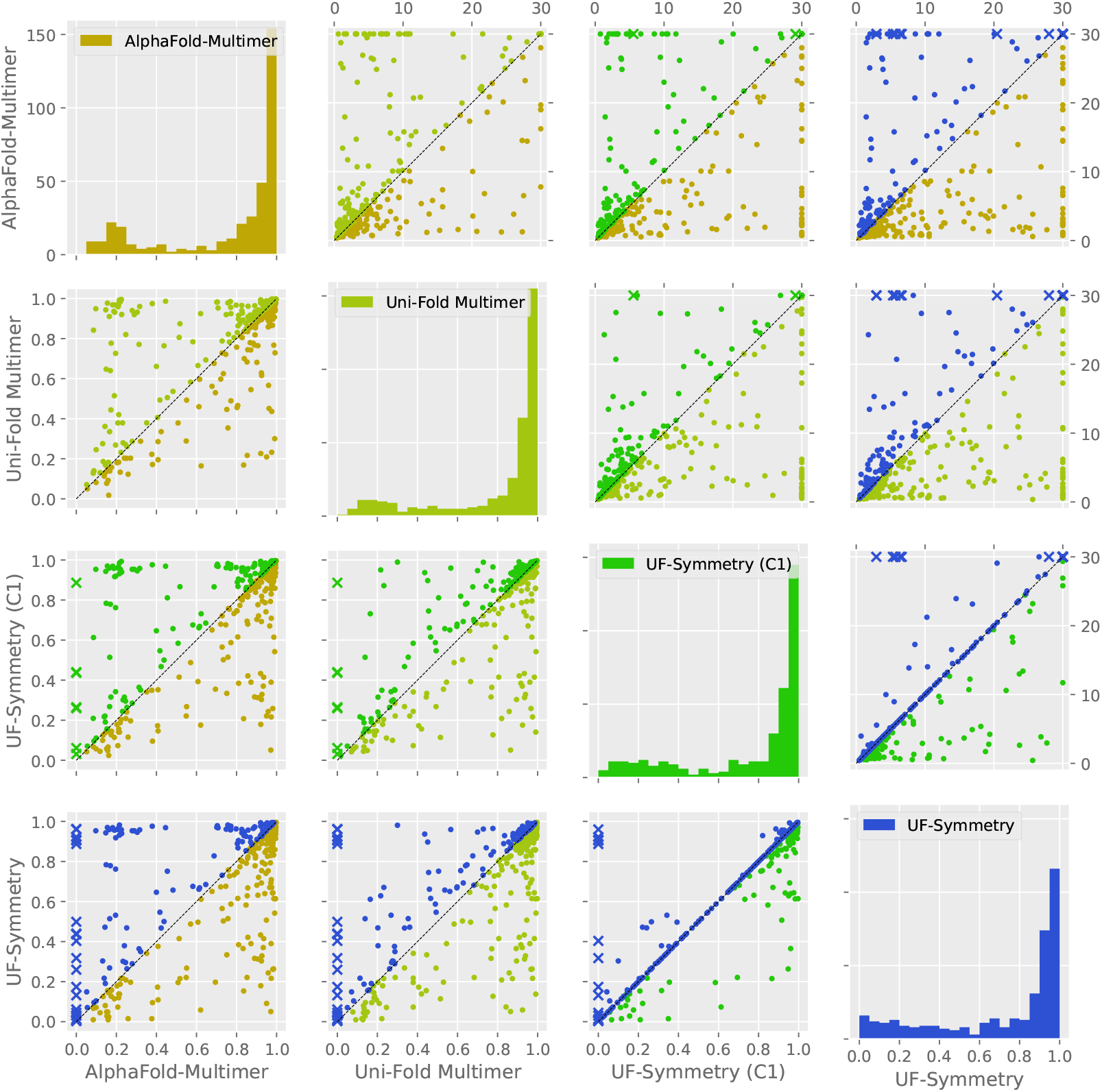
Scatter matrix of accuracies of UF-Symmetry and baselines. The lower triangle stores the TM-Score scatter plots, the diagonal stores the histograms of TM-Score, and the upper triangle stores the C_*α*_-RMSD_30_ scatter plots. Dots in different colors indicate that the corresponding model is better (blue for UF-Symmetry). Crosses indicate that the other model failed to predict the structure.

#### Case Study

Figure 6 presents one of the largest targets in the test set, which is a homo 12-mer structure (PDB-ID: 7SYA) in the C12 symmetry group. We show both the assembly and the AU structures predicted by UF-Symmetry, aligned with the experimentally solved ones. As a total of 8,460 residues is deposited in the assembly, AlphaFold-Multimer, Uni-Fold Multimer and UF-Symmetry (C1) failed to predict for this target. UF-Symmetry successfully predicted the structure of the AU, and correctly assembled the complex. The model inference time for the assembly is approximately 80 seconds on an NVIDIA A100 GPU.

**Figure 6:**
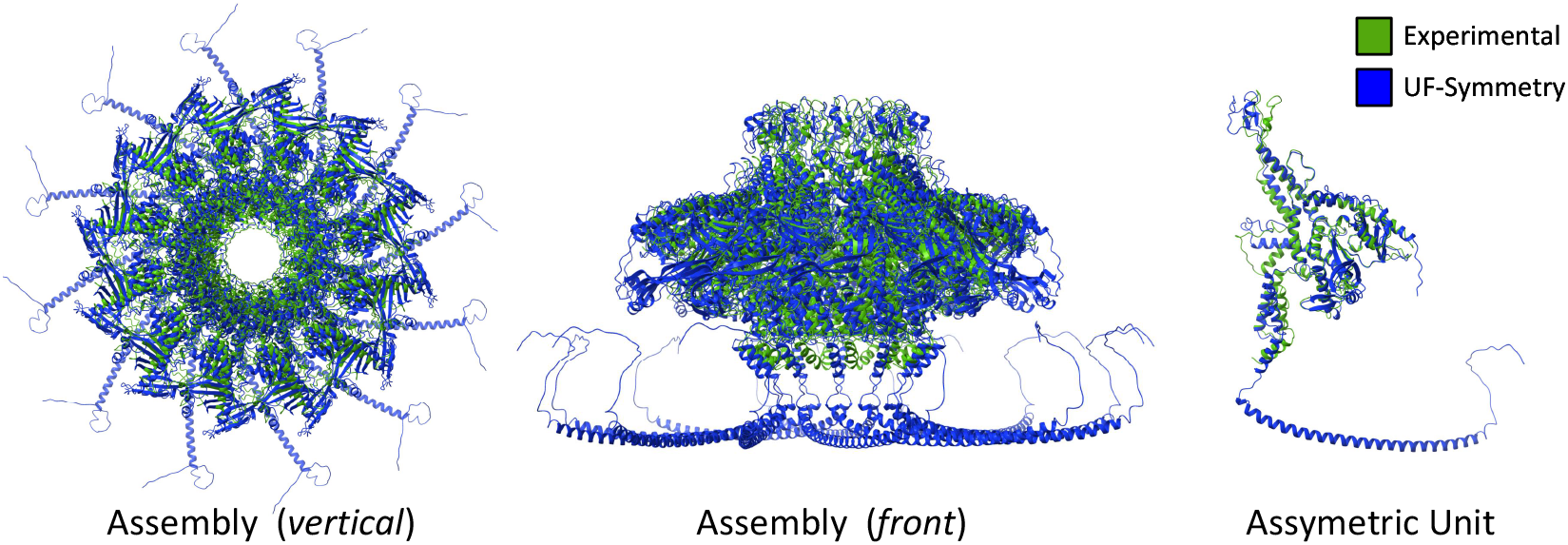
Predicted results of a *homo 12-mer* structure (PDB-ID: 7SYA) by UF-Symmetry. The total number of residues is 8,460. The TM-Score of the prediction is 0.922. The long helix tails in the prediction are not experimentally solved.

## 5 Conclusion and Future Work

In this work, we propose UF-Symmetry a fast and accurate solution to predicting the structures of large symmetric protein oligomers. UF-Symmetry reduces the need of copying the sequences of asymmetric units, thus achieving great advantages over model scalability and efficiency, yet with a slight drop of average accuracy.

UF-Symmetry is currently an on-going work, with straight-forward upcoming extensions in i) design of model confidence; ii) training on dihedral and cubic symmetry groups; iii) various optimizations inherited from Uni-Fold, and iv) larger evaluation sets. In addition, UF-Symmetry currently relies on pre-specified symmetry groups, and hence another promising extension is to automatically extract the possible symmetry groups given the AU sequences.

## A Statistics of Protein Symmetries in PDB

We made statistics on the 181,727 protein structures in the Protein Data Bank (PDB) [6] released before Janurary 16th, 2022. We used the symmetry symbols annotated by the RCSB website^7^. Approximately 54% PDB structures are multimeric structures, among which approximately 72% are symmetric. Table 2 shows the detailed statistics.

**Table 2:** Statistics of symmetries of PDB structures

| Type | Symmetry | Number of structures |
| --- | --- | --- |
| Monomers | – | 83,392 |
| Multimers | Asymmetric | 27,470 |
|  | C2 | 45,496 |
|  | C3 | 6,037 |
|  | C4 | 1,736 |
|  | C5 | 893 |
|  | C6 | 639 |
|  | Larger cyclic | 587 |
|  | D2 | 8,571 |
|  | D3 | 2,577 |
|  | D4 | 815 |
|  | D5 | 302 |
|  | D6 | 191 |
|  | Larger dihedral | 239 |
|  | Icosahedral | 1,182 |
|  | Octahedral | 544 |
|  | Tetrahedral | 475 |
|  | Helical | 581 |
|  | **All** | 98,335 |
| Total | – | 181,727 |

## B Standard Symmetry Operations of Cubic Symmetries

The group of the standard operations of the icosahedral symmetry is the multiplication of a 5-fold rotation group of the *z* axis ((0, 0, 1)), a 3-fold rotation group of axis 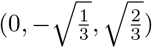, a 2-fold rotation group of axis 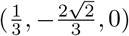, and a 2-fold rotation group of 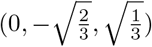.

The group of the standard operations of the octahedral symmetry is the multiplication of a 4-fold rotation group of the *z* axis ((0, 0, 1)), a 3-fold rotation group of axis 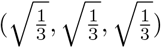, and a 2-fold rotation group of the *x* axis (1, 0, 0).

The group of the standard operations of the tetrahedral symmetry is the multiplication of a 3-fold rotation group of axis 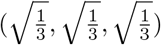, a 2-fold rotation group of the *y* axis ((0, 1, 0), and a 2-fold rotation group of the *x* axis (1, 0, 0).

## C Proofs of Theorem 1 and Equations (7), (8) and (9)

To prove Theorem 1, we first introduce Lemma 1:

### Lemma 1

*For any finite group* 𝒢 *with* | 𝒢| = *K and any real-valued function* ℱ : 𝒢 → ℝ,

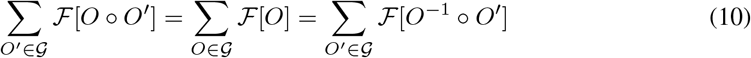

The proof of Lemma 1 is simple by noticing that for any *O* ∈ 𝒢, we have *O*^−1^ ∈ 𝒢, and applying *O* on all elements in 𝒢 generates a permutation of 𝒢, i.e. *O* ○ 𝒢 = *O*^−1^ ○ 𝒢 = 𝒢.

Assuming the assembly 𝒮 conforms to the symmetry group 𝒢, Theorem 1 can then be proved as

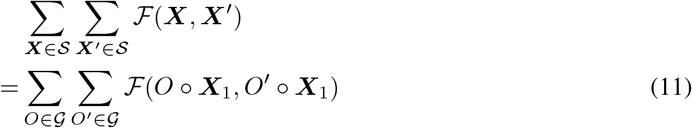

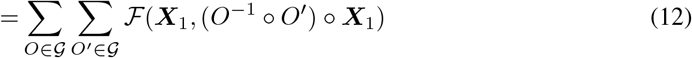

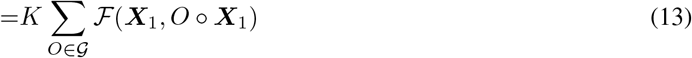

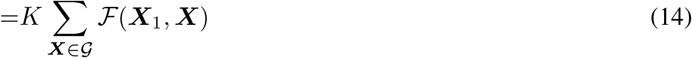

where Equation (11) holds as 𝒮 conforms to symmetry group 𝒢, Equation (12) holds under the condition that ℱ is invariant under 3D transformations, and Equation (13) is derived from Lemma 1.

The equivalency of FAPE loss follows

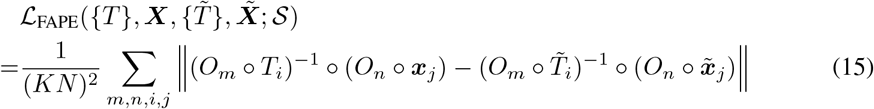

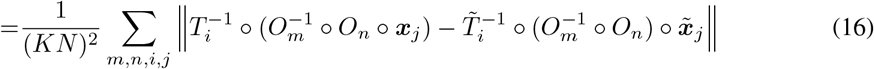

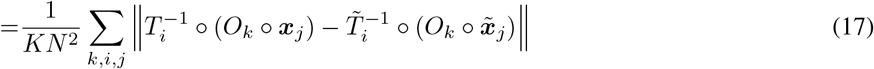

The equivalency of *between-AU* clash loss follows

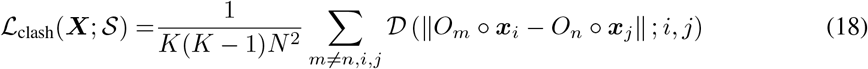

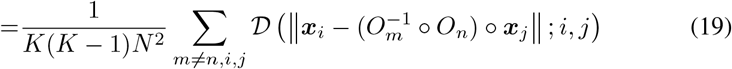

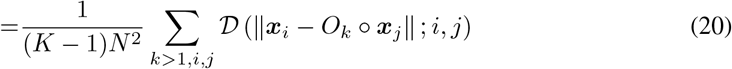

where 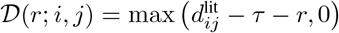 is the kernel function of the loss.

The equivalency of *between-AU* chain centre-of-mass follows

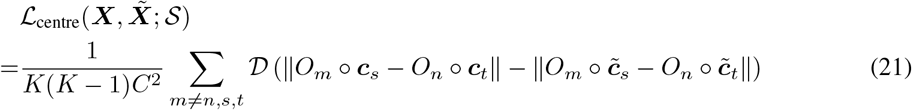

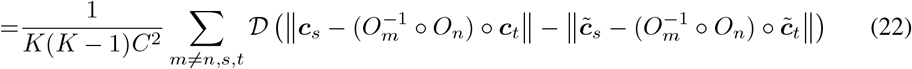

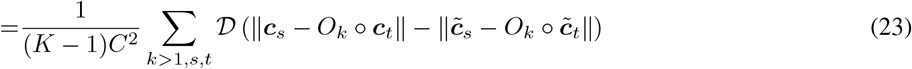

where 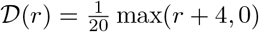.

## Footnotes

2 We use the term *oligomers* to refer to protein complexes of two or more repeating subunits.

3 Results may have varied now.

4 We use the term *assembly* for a protein complex that undertake biological functions as a whole, the term *asymmetric units* (AUs) for its repeating components. Notably, AUs may contain one or more chains.

5 https://www.rcsb.org/.

6 https://github.com/dptech-corp/Uni-Fold. We developed UF-Symmetry and conducted experi-ments on an early commit of Uni-Fold (commit-ID: b7461325). As the repository lately introduced plenty of optimizations, the capacity and efficiency results may have varied.

7 https://www.rcsb.org/

